# Effect of pressure insole sampling frequency on peak force accuracy during running

**DOI:** 10.1101/2022.05.18.492523

**Authors:** L.J. Elstub, L.M. Grohowski, D.N. Wolf, M.K. Owen, B. Noehren, K.E. Zelik

## Abstract

Pressure sensing insoles enable us to estimate forces under the feet during activities such as running, which can provide valuable insight into human movement. Pressure insoles also afford the opportunity to collect more data in more representative environments than can be achieved in laboratory studies. One key challenge with real-world use of pressure insoles is limited battery life which restricts the amount of data that can be collected on a single charge. Reducing sampling frequency is one way to prolong battery life, at the cost of decreased measurement accuracy, but this trade-off has not been quantified, which hinders decision-making by researchers and developers. Therefore, we characterized the effect of decreasing sampling frequency on peak force estimates from pressure insoles (Novel Pedar, 100 Hz) across a range of running speeds and slopes. Data were downsampled to 50, 33, 25, 20, 16 and 10 Hz. Force peaks were extracted due to their importance in biomechanical algorithms trained to estimate musculoskeletal forces and were compared with the reference sampling frequency of 100 Hz to compute relative errors. Peak force errors increased exponentially from 0.7% (50 Hz) to 9% (10 Hz). However, peak force errors were <3% for all sampling frequencies down to 20 Hz. For some pressure insoles, sampling rate is inversely proportional to battery life. Therefore, these findings suggest that battery life could be increased up to 5x at the expense of 3% errors. These results are encouraging for researchers aiming to deploy pressure insoles for remote monitoring or in longitudinal studies.

## Introduction

Pressure sensing insoles enable us to measure forces under the feet during activities such as running, which can provide valuable insight into human movement and clinical care (Chuckpaiwong et al., 2008; Dixon, 2008; Hullfish and Baxter, 2020; Mann et al., 2016). For instance, we have previously shown that peak force under the foot, which can be estimated from a pressure insole, is an important input for biomechanical algorithms trained to estimate forces on and understand injury risk to musculoskeletal structures inside the body (Matijevich et al., 2020). Pressure insoles can also be used in real-world data collections outside the laboratory, to collect more data (e.g., more strides), in more realistic and representative environments, and over longer periods than can be achieved in laboratory studies. This portable foot pressure measurement capability unlocks new possibilities for longitudinal studies and real-world monitoring which can empower us to better understand human movement, behaviour, performance, injury mechanisms, and associated risk factors.

However, a practical challenge of pressure insoles is battery power which limits the amount of data that can be collected on a single charge. This limitation is not a problem for short data collections (e.g., in a laboratory setting) but often becomes an issue when trying to collect data for longer durations (e.g., an entire day, multiple days). One parameter that affects the battery life is the size (capacity) of the battery, but there are constraints on this size to maintain lightweight and low-profile devices worn in/on a shoe. Another parameter that affects battery life is the sampling frequency, the number of samples recorded by the device each second. A simple way to conserve battery life is to reduce sampling frequency; however, if reduced too much this can degrade pressure measurement accuracy and the estimation of important metrics (e.g., peak forces). Although the Nyquist frequency provides a theoretical lower bound on the sampling frequency needed for an activity, it is generally not practical for informing appropriate sampling frequency in clinical or real-world environments. Empirical studies are often needed to characterize how the choice of sampling frequency affects measurement accuracy (e.g., Renner et al., 2022) and this information is critical for researchers and device designers to make informed decisions.

Research-grade pressure insoles often sample data on the order of 100 - 200 Hz which is sufficient for biomechanical analysis of most activities (e.g., walking, running, jumping, Peebles et al., 2018; Renner et al., 2022, 2019) and sufficient for most laboratory studies (which generally last a few hours or less). However, when using pressure insoles for continuous, all-day, or multi-day monitoring in the real-world, battery power can become a significant barrier. Additionally, high sampling frequencies require large memory capacity and longer data processing time. Presently, there is little empirical data in the scientific literature to help elucidate the degree to which lower sampling frequencies degrade estimates of key biomechanical metrics extracted from pressure insoles. We encountered this barrier ourselves when trying to assess what range of sampling frequencies would be acceptable for monitoring running in the real-world, given our desire to capture the active force peak under the foot each step as a key input to musculoskeletal force estimation algorithms we previously developed (Matijevich et al., 2020). To overcome this barrier, the purpose of this study was to characterise the effect of decreasing sampling frequency on the accuracy of peak force estimates from pressure insoles during running.

## Methods

This study was performed as a secondary analysis of data collected in a previous study (Elstub et al., *under review*). All participants were actively running a minimum of ten miles per week and were not experiencing any injuries at the time of data collection or six months prior. The data collection was approved by the Vanderbilt University Institutional Review Board and all participants gave written informed consent before participating.

We collected data from nine recreational runners (6 males [height: 1.84 ± 0.07 m, mass: 82.4 ± 7.9 kg], 3 females [height: 1.72 ± 0.08 m, mass: 59.3 ± 4.4]) while running on a treadmill during 40 trials. For each trial, the participants ran for 30 seconds, and the final 10 left steps of a trial were chosen for analysis. To ensure a variety of running conditions were analysed in this study, the participants ran across a range of slopes (0°, 3°, 6°, −3°, and −6°) and speeds (2.68 m/s, 2.95 m/s, and 3.31 m/s, which correspond with approximately 8–10 minute/mile paces). All participants in the study wore ASICS Gel Cumulus 20 trainers to reduce confounding variables (e.g., pressure insole fit issues) arising from the use of different running shoes.

The normal plantar force of each foot during each step was estimated by Novel pressure sensing insoles (Pedar, Munich, Germany) at a maximum sampling frequency of 100 Hz. This was taken to be the reference force measurement. To quantify the effect of different sampling rates, we downsampled the reference data to 50, 33.3 (referred to as 33 Hz), 25, 20, 16.7 (referred to as 16 Hz), and 10 Hz using the function `downsample` in MATLAB R2019b (Natick, MA). The downsample function was chosen to maintain the integrity of the signal and prevent filtering methods from biasing the calculation of peak normal force. In other words, this simulates data that would have resulted if only every *n*^th^ sample had been collected by the pressure insole (e.g., 20 Hz is every fifth sample). For each step, the active force peak was computed separately for each sampling rate. We focused on active peak force from the pressure insoles because we previously found it to be the most important input signal for a wearable sensor system that was trained to estimate peak tibial bone force (Matijevich et al., 2020).

### Analysis

For each sampling rate less than 100 Hz, we plotted the normal force waveform (Figure 1) and then computed the percent error in active peak force on a step-by-step basis compared with the 100 Hz condition. We calculated the mean percent error (referred to as *average error* throughout) across all 10 steps during a trial for each participant, then normalized it by the 100 Hz active peak force to report errors as percentages. For this study, the data presented are for the left foot only as we expect the same conclusions to hold for both sides. We did not perform statistical analysis on the effect of sampling frequency because the purpose of the study was not to determine if peak force errors increase with a lower sampling frequency, nor did we have a specific hypothesis related to sampling frequency. Rather, the purpose of this study was to characterise the effect of decreasing sampling frequency on the accuracy of peak force estimates from pressure insoles to provide quantitative data to inform decision-making. As such, researchers and wearable device developers should use these results to consider what level of accuracy is necessary, or acceptable, for their specific purpose and choose a sampling frequency accordingly.

**Figure 1.**
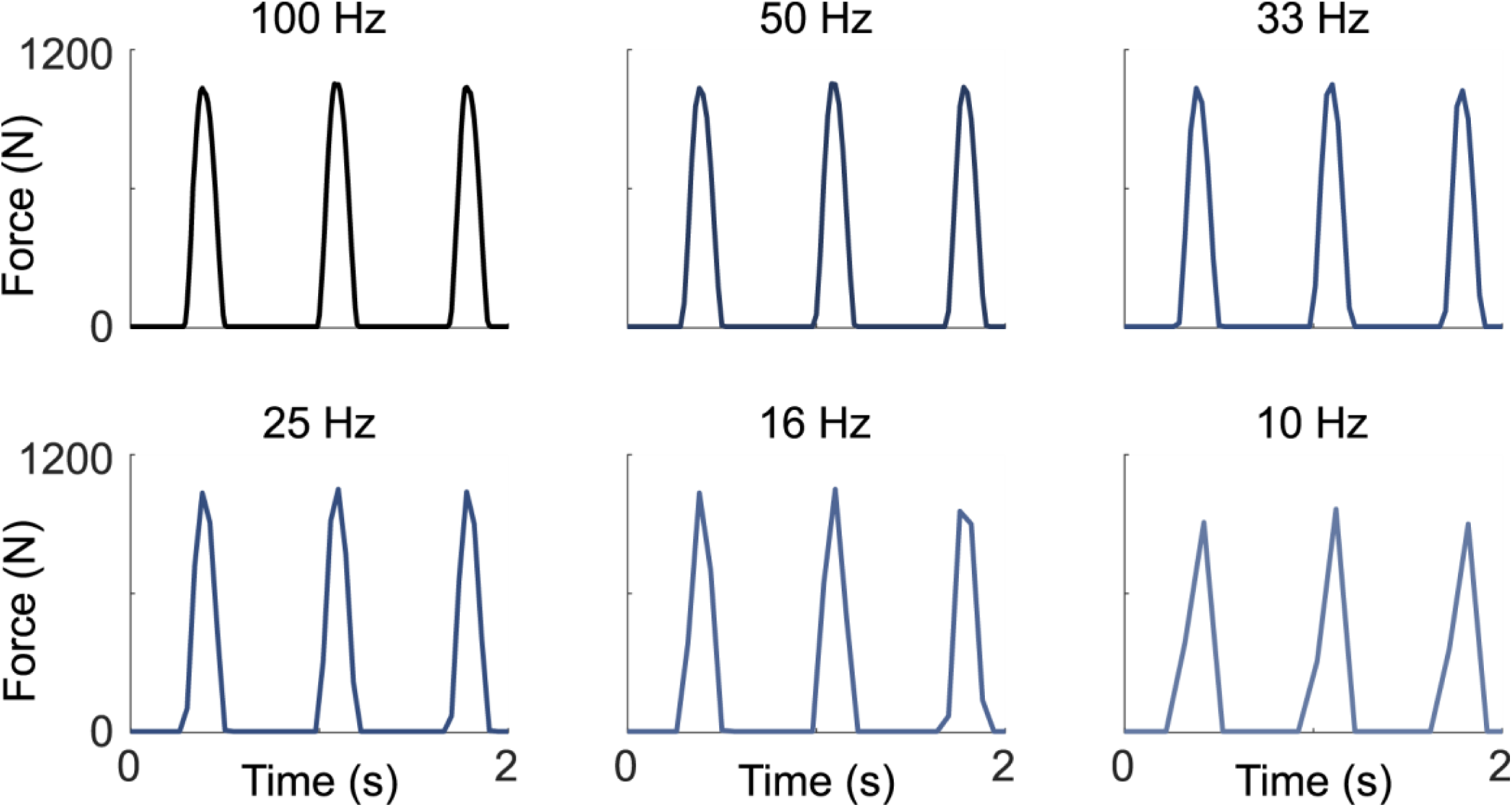
Representative force-time curves for one participant running at 3.31 m/s on a zero degree slope at different sampling frequencies

## Results

Errors in active peak force increased as sampling frequency decreased from the reference value of 100 Hz (Figure 2). Across all running conditions, average errors of 0.7%, 1.3%, 1.8%, 2.8%, 3.7% and 8.7% were observed at 50, 33, 25, 20, 16 and 10 Hz, respectively. As sampling frequency decreased from 100 to 10 Hz, we also observed a decrease in mean active peak force for all participants (Figure 1, Table 1). Across all participants, this resulted in a mean 32 N decrease (∼5% body weight for the average mass of 73 kg in this study) in mean peak normal force at 20 Hz compared with the 100 Hz reference condition, and a 116 N decrease at 10 Hz (∼16% body weight for a 73 kg individual).

**Table 1.** Mean active peak force (N) across all trials (speeds and slopes) for each participant at different sampling frequencies. Each enumerated column is one participant.

| Sampling Frequency (Hz) | Peak Force (N) | | | | | | | | | Mean ± std |
| --- | --- | --- | --- | --- | --- | --- | --- | --- | --- | --- |
|  | 1 | 2 | 3 | 4 | 5 | 6 | 7 | 8 | 9 |  |
| 10 | 1135 | 1015 | 1255 | 787 | 756 | 1427 | 1811 | 1387 | 697 | **1141 ± 348** |
| 16 | 1231 | 1077 | 1318 | 819 | 810 | 1485 | 1951 | 1468 | 745 | **1212 ± 372** |
| 20 | 1250 | 1084 | 1330 | 828 | 822 | 1502 | 1973 | 1479 | 753 | **1225 ± 376** |
| 25 | 1265 | 1093 | 1340 | 830 | 834 | 1535 | 2004 | 1492 | 762 | **1239 ± 384** |
| 30 | 1273 | 1101 | 1347 | 837 | 841 | 1526 | 2021 | 1502 | 767 | **1246 ± 385** |
| 50 | 1282 | 1104 | 1353 | 840 | 848 | 1551 | 2035 | 1511 | 771 | **1255 ± 389** |
| 100 | 1287 | 1102 | 1356 | 843 | 836 | 1555 | 2043 | 1516 | 774 | **1257 ± 392** |

**Figure 2.**
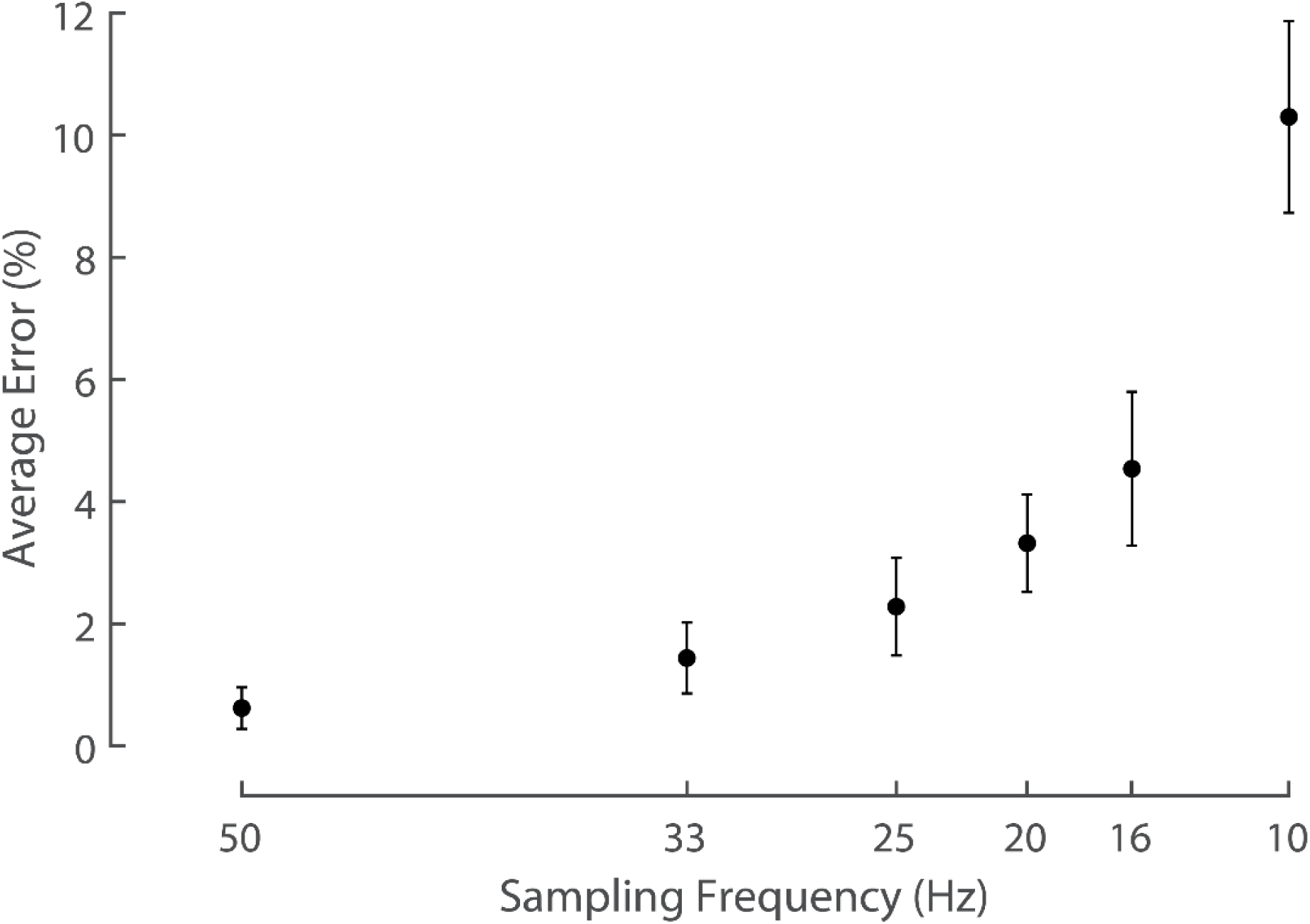
Average error in mean active peak force across all participants and trials for each sampling frequency compared with the reference sampling frequency of 100 Hz. Error bars represent standard deviation between participants.

Errors in peak force due to downsampling were consistent across speeds and slopes, and this is exemplified for the 20 Hz sampling frequency in Table 2.

**Table 2.** Average error (%) of mean peak force across speeds and slopes for the 20 Hz condition compared with 100 Hz. The negligible trends across speed and slope presented here are representative of trends observed at other sampling frequencies.

| Slope<br>(degrees) | Speed (m/s) |  |  |
| --- | --- | --- | --- |
|  | 2.68 | 2.95 | 3.31 |
| -6 | 3.8% ± 1.6% | 3.4% ± 1.6% | 4.3% ± 2.2% |
| -3 | 2.8% ± 0.7% | 2.5% ± 1.1% | 3.3% ± 1.3% |
| 0 | 2.7% ± 1.4% | 2.6% ± 1.2% | 3.0% ± 1.3% |
| 3 | 2.7% ± 1.1% | 2.2% ± 0.8% | 2.5% ± 1.1% |
| 6 | 2.5% ± 0.7% | 2.8% ± 1.1% | 2.6% ± 0.7% |

## Discussion

We quantified the effect of different sampling frequencies on estimating active peak force from pressure sensing insoles. Peak force errors increased exponentially from 0.7% error at 50 Hz to 9% error at 10 Hz, relative to the 100 Hz reference sampling frequency. However, average errors in peak force across speeds and slopes were <3% for all sampling frequencies down to and including 20 Hz. These results are highly encouraging for researchers and developers aiming to deploy pressure insoles in longitudinal or long-duration studies, or for real-world monitoring.

Results from the present study demonstrate that it is possible to capture accurate peak force estimates with pressure insoles without high sampling rates commonly used in lab studies. Previous research has recommended pressure insole sampling frequencies in the range of 100 Hz and above, based on the accuracy of peak and mean force, time to peak force, and peak rate of force development during walking, running, and countermovement jumps (Hori et al., 2009; Renner et al., 2022). However, these lab-focused recommendations did not incorporate or consider other aspects of device usability, such as limited battery power or data storage space, that are necessary to address for translation to certain real-world applications.

In this study, we found that sampling frequency can be reduced by a factor of 5 (from 100 Hz to 20 Hz) while only introducing ∼3% error in peak force. For some pressure insoles, the sampling rate is inversely proportional to battery life (based on conversations with insole manufacturers). For these specific cases, battery life could be increased by up to 5x at the expense of 3% peak force errors. For many real-world applications, this is a phenomenally good trade-off. Additionally, lower sampling rates produce smaller file sizes which increase the capacity for data storage in onboard memory. It is recommended that researchers and wearable technology developers consider device needs and trade-offs holistically, and then determine what level of error is acceptable for their application.

There are limitations to the current study. We only examined one brand and model of pressure insole (Novel Pedar). However, we expect the main conclusions are generalisable across manufacturers and models, and not heavily dependent on the number or orientation of pressure sensors in the insole. In addition, we tested a limited number of speeds. Whilst we did not observe a change in peak force accuracy due to speed (Table 2), we expect this might change at much faster speeds due to the shorter ground contact times that occur (Weyand et al., 2000). Therefore, we expect the results here to be informative and applicable for long-distance running at paces similar to those studied here, and for ambulation tasks with similar or longer ground contact times (e.g., walking, activities of daily living). However, these findings may not extrapolate to more impulsive sprinting or jumping activities. Finally, the results presented apply specifically to our chosen metric of active peak force. We expect higher sampling frequencies are necessary to accurately capture other running metrics (e.g., loading rate, force impulse; Renner et al., 2022) which change at lower sampling frequencies (Fig. 1).

## Conclusion

We estimated active peak force during running with pressure sensing insoles and found that average errors were <3% for all sampling frequencies down to and including 20 Hz, relative to nominal 100 Hz sampling. These findings can inform decision-making related to pressure insole sampling rate for real-world applications or longitudinal studies, in which it is necessary to balance trade-offs between measurement accuracy and other factors such as battery life or onboard data storage capacity.

